# A dual role for CRTH2 in acute lung injury

**DOI:** 10.1101/2022.05.29.493897

**Authors:** Shreya Bhattacharya, Nick Ristic, Avi J. Cohen, Derek Tsang, Meredith S. Gwin, Rebecca Howell, Grant M. Young, Eric Jung, Charles S. Dela Cruz, Samir Gautam

**Affiliations:** Division of Pulmonary, Critical Care, and Sleep Medicine, Department of Internal Medicine, Yale School of Medicine – New Haven, CT/US; Department of Chemistry, Yale University – New Haven, CT/US; Department of Microbial Pathogenesis, Yale School of Medicine – New Haven, CT/US

**Keywords:** Neutrophils, CRTH2, acute lung injury, PGD2, type 2 immunity, murine ARDS model

## Abstract

Acute respiratory distress syndrome (ARDS) is a life-threatening clinical condition defined by rapid-onset respiratory failure following acute lung injury (ALI). The high mortality rate and rising incidence of ARDS due to COVID-19 make it an important research priority. Here we sought to investigate the role of chemoattractant receptor-homologous molecule expressed on Th2 cells (CRTH2) in ARDS. CRTH2 is a G protein-coupled receptor best studied in the context of type 2 immunity, but it also exerts effects on neutrophilic inflammation. To evaluate its role in mouse models of ARDS, we first examined its expression pattern on murine neutrophils. We found it is expressed on neutrophils, but only after extravasation into the lung. Next, we showed that CRTH2 expression on extravasated lung neutrophils promotes cell survival, as genetic deletion of CRTH2 and pharmacologic inhibition of CRTH2 using fevipiprant both led to increased apoptosis in vitro. We then evaluated the role of CRTH2 in vivo using a murine model of LPS-induced ALI. In line with the pro-inflammatory effects of CRTH2 in vitro, we observed improvement of lung injury in CRTH2-deficient mice in terms of vascular leak, weight loss and survival after LPS administration. However, neutrophilic inflammation was elevated, not suppressed in the CRTH2 KO. This finding indicated a second mechanism offsetting the pro-survival effect of CRTH2 on neutrophils. Bulk RNAseq of lung tissue indicated impairments in type 2 immune signaling in the CRTH2 KO, and qPCR and ELISA confirmed downregulation of IL-4, which is known to suppress neutrophilic inflammation. Thus, CRTH2 may play a dual role in ALI, directly promoting neutrophil cell survival, but indirectly suppressing neutrophil effector function via IL-4.

## Introduction

Acute respiratory distress syndrome (ARDS) is a life-threatening clinical condition defined by rapid onset respiratory failure following acute lung injury (ALI).^1^ Due to the COVID-19 pandemic, ARDS has emerged as a leading cause of morbidity and mortality worldwide.^2^ Yet despite its growing clinical burden, no effective disease-modifying therapies have been identified, outside of steroids for pneumonia-associated lung injury.^1^ Thus, investigations into ARDS pathogenesis and treatment represent an important ongoing research priority.

Chemoattractant receptor-homologous molecule expressed on Th2 cells (CRTH2, also called DP2, CD294, GPR44, and PTGDR2) is a seven-membrane spanning G-protein coupled receptor (GPCR) expressed on type 2 immune cells such as eosinophils, mast cells, Th2 cells, and type 2 innate lymphoid cells (ILC2s).^3–5^ Upon activation with its endogenous ligand, prostaglandin D_2_ (PGD_2_), CRTH2 signals through G_i_α, leading to increases in cytosolic calcium and reduction in cAMP levels.^4^ In eosinophils, CRTH2 agonism promotes inflammatory responses including degranulation and chemotaxis.^3^ CRTH2 ligation similarly activates ILCs and Th2 cells, leading to chemotaxis, suppression of apoptosis, and generation of type 2 cytokines including IL-4, IL-5, IL-9, and IL-13.^6–9^

Given its key role in type 2 inflammation, CRTH2 has been investigated as a therapeutic target in allergic diseases such as asthma. Numerous small molecule antagonists have been developed for this purpose, as best exemplified by fevipiprant, which advanced to late-stage clinical trials.^10^ Phase II studies in asthma showed promising results, as fevipiprant suppressed peripheral and airway eosinophilia, reduced airway remodeling, improved quality of life, and increased lung function.^11–14^ In a follow-up phase III clinical trial, CRTH2 blockade proved safe and efficacious as it improved lung function and asthma control,^15^ but further development was halted as it failed to reduce exacerbation frequency.^10^

The function of CRTH2 outside of type 2 immunity is less understood, but several studies have indicated a role in neutrophilic inflammation.^16^ For instance, CRTH2 inhibition reduced neutrophil recruitment to the lung in a lipopolysaccharide (LPS)-induced model of ALI.^17^ Consistent results were observed upon pharmacologic blockade of CRTH2 in murine smoking models.^18,19^ Similarly, Jandl et al demonstrated that activation of CRTH2 with the isoform-specific agonist, DK-PGD2, exacerbated neutrophilic inflammation during ALI.^20^ Outside of the lung, CRTH2 appears to play a similar pro-neutrophilic role, as shown in murine models of cutaneous inflammation.^21,22^

Thus, we hypothesized that CRTH2 inhibition might ameliorate neutrophil-mediated tissue damage during ALI, and therefore represent a novel therapeutic strategy for ARDS. To evaluate this hypothesis, we studied the expression and function of CRTH2 in murine neutrophils isolated from the lung. We then utilized a model of LPS-induced ALI in CRTH2 knockout mice to investigate the function of CRTH2 in vivo.

## Methods

### Murine model of acute lung injury

Male and female wild-type and CRTH2 KO C57BL/6 mice (Jackson Laboratories) were bred at Yale University under specific pathogen-free conditions. All animal studies were performed in accordance with Institutional Animal Care and Use Committee (IACUC) approved protocols at Yale University. 8-to 12-week-old age- and gender-matched mice were used for experiments after randomizing to control for litter effects. Intrapulmonary instillation of *Pseudomonas aeruginosa*-derived LPS (Sigma, L9143) via oropharyngeal (OP) aspiration, performance of BAL, measurement of cell counts, and quantification of neutrophil elastase activity were performed as described previously.^23,24^ To induce ALI, LPS was delivered according to weight-based dosing (0.25-10 μg LPS per gram of mouse body weight) and mice were sacrificed at defined endpoints.

### Flow cytometry

To determine neutrophil counts, total white blood cells (WBCs) were counted on a benchtop hematology analyzer (AcT Diff, Beckman Coulter) and BAL cells were stained with antibodies against Ly6G (Biolegend) and CD45 (Biolegend) prior to evaluation by flow cytometry, as previously described.^23^ To quantify CRTH2 surface expression, blood and BAL cells were stained for Ly6G, F4/80 (Biolegend), and CRTH2 (Thermo). An isotype matched antibody was used to control for non-specific CRTH2 staining.

### In vitro assay for neutrophil apoptosis

Neutrophils were collected from mice via BAL 12-24 hours after intrapulmonary delivery of 5 μg of LPS, isolated to >95% purity using the MojoSort™ Mouse Neutrophil Isolation Kit (480058, BioLegend) following the manufacturer’s instructions. Cells were washed twice in PBS, centrifuged at 2600 RCF for 6 minutes, and resuspended in Dulbecco’s Modified Eagle Medium/F12 without phenol red (Thermo). Wells were loaded with the neutrophil suspension (200,000 total cells), and indicated stimuli were then added together with the fluorogenic caspase 3/7 substrate CellEvent (C10432, Thermo), at a final concentration of 2 μM. Kinetics of apoptosis was quantified with a BioTek Cytation 3 plate reader by measuring fluorescence at 10-minute intervals over 12h (excitation 502 nm; emission 530 nm).

### Transcriptomic analysis

Whole lungs were snap-frozen in 500 μL TRIzol buffer (Invitrogen) and homogenized via sonication at 20 kHz for 1 min before RNA isolation was performed using a RNeasy Mini Kit (Qiagen). For qPCR, the iScript cDNA Synthesis Kit (Bio-Rad) was utilized to generate cDNA; 18S rRNA was used as the reference gene. For bulk RNA-seq, quality assessment and cDNA library preparation were performed by the Yale Center for Genomic Analysis prior to rRNA depletion and Illumina-based NovaSeq S2 sequencing using a read length of 2x100 and 35M reads/sample.

### Statistical analysis

Statistical analyses were performed using Prism 8 (Graphpad Software Inc). Results are presented as mean ± standard error of the mean (SEM). Data were analyzed using Student’s t-test or one-way ANOVA as appropriate. Data shown are representative of multiple independent repetitions of the experiment. Statistical significance throughout was defined as *P* < 0.05.

RNA sequencing (RNAseq) analysis was conducted using the *edgeR* package in the R programming language (V.3.6.0, Vienna, Austria), and potential differentially expressed genes (“candidate DEGs”) were identified by selecting genes with both a raw p-value of < 0.05 and consistent direction of regulation within sample groups (allowing for no more than one outlier per gene) after TPM normalization and scaling. We performed KEGG pathway analysis and gene ontology (GO) analysis using the Enrichr analysis tool as described by Chen et al.^25^

## Results

### Flow cytometry demonstrates cell-surface expression of CRTH2 on neutrophils

To assess whether CRTH2 has direct functional effects on neutrophils, we first tested for cellular expression of the receptor. Using flow cytometry we found that CRTH2 was expressed on the surface of F4/80+ cells in the blood (such as monocytes and eosinophils), consistent with prior reports (Fig. 1).^3^ We also observed that it was absent from circulating neutrophils, consistent with prior reports.^26^ However, after recruitment into the lung, extravasated neutrophils collected from the bronchoalveolar lavage (BAL) showed significant staining for CRTH2, indicating compartment-specific receptor expression. Notably, CRTH2+ neutrophils were only detected in the lung during the first 24 hours after LPS administration, indicating that they may represent an inflammatory subtype recruited early during ALI.

**Figure 1:**
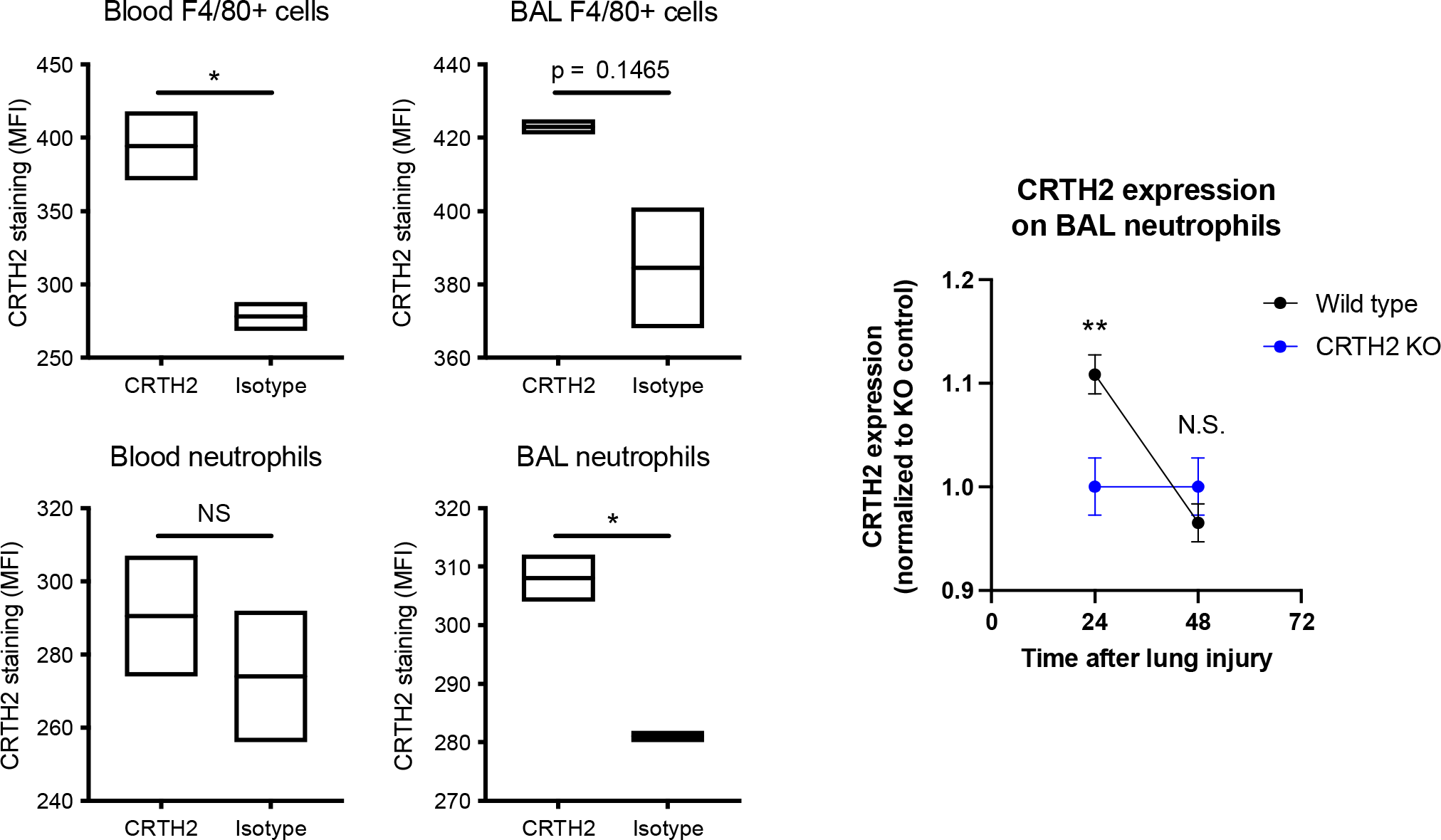
CRTH2 is expressed on neutrophils after extravasation into the lung. Blood and BAL were collected from mice with ALI. Cells were stained for Ly6G (indicating neutrophils), F4/80 (indicating monocytes, macrophages, and eosinophils), and CRTH2. An isotype antibody for CRTH2 was used to control for non-specific binding. F4/80^+^ cells were CRTH2 positive. Neutrophils only demonstrated CRTH2 expression after extravasation into the lung during the first 24h.

### CRTH2 mediates pro-inflammatory responses in neutrophils in vitro

BAL neutrophils were then tested for functional responses to CRTH2 agonists. To do so we utilized DK-PGD2, which binds selectively to CRTH2 and not DP1 (a closely related prostanoid receptor also capable of binding PGD2).^4^ In a microplate-based kinetic fluorescence assay measuring caspase 3/7 activity, BAL neutrophils treated with DK-PGD2 failed to show any differences in apoptosis (data not shown). However, treatment with the CRTH2-specific antagonist, fevipiprant, demonstrated increased apoptosis in a CRTH2-dependent manner (Fig. 2). In addition, we found that CRTH2 KO neutrophils showed an intrinsic defect in survival both at baseline and upon stimulation of apoptosis with staurosporine.

**Figure 2:**
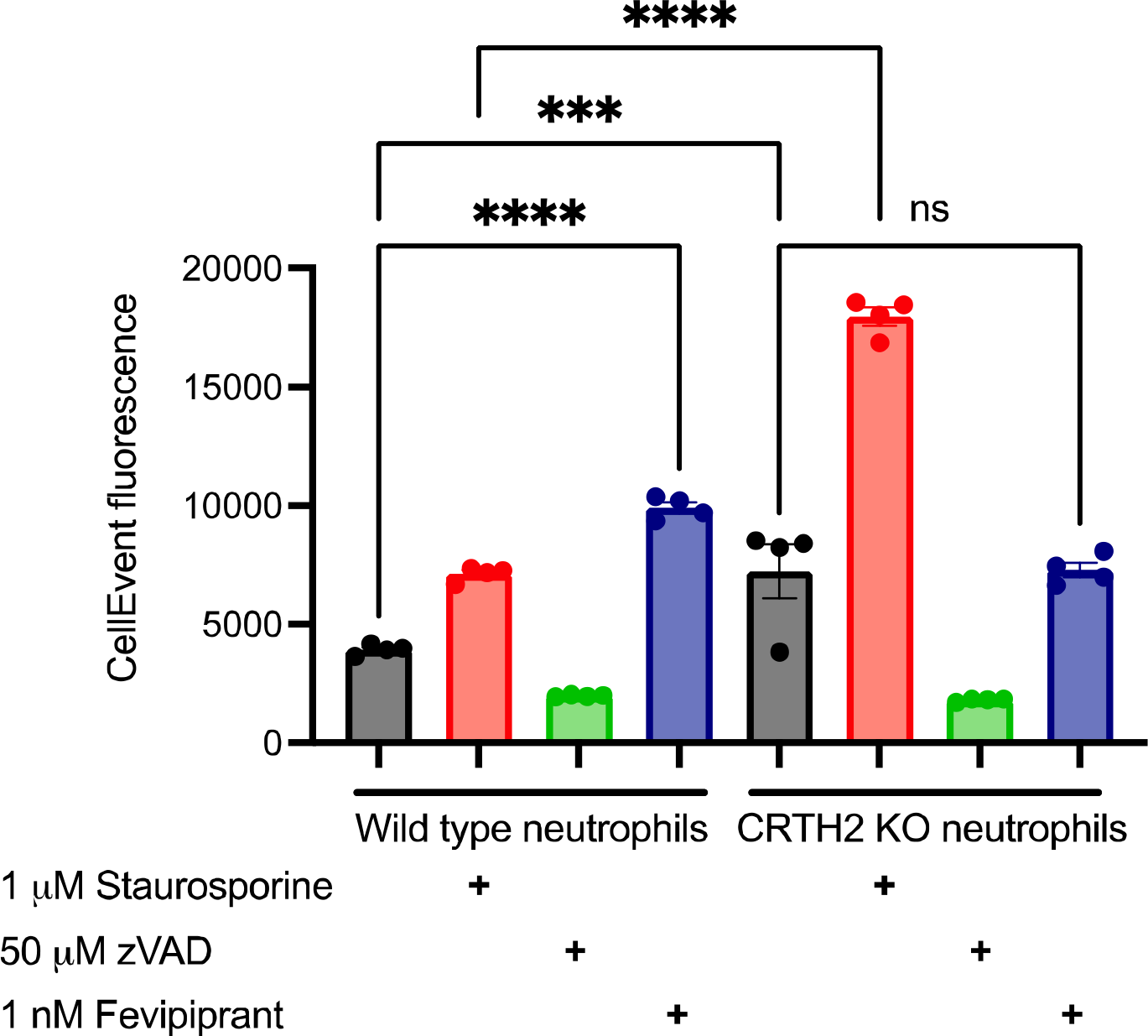
CRTH2 mediates pro-inflammatory responses in lung neutrophils. 5 μg of LPS was delivered to the lung induce lung neutrophilia. BAL was collected and isolated neutrophils were incubated with CellEvent, a fluorescent sensor for apoptosis. The CRTH2 antagonist fevipiprant was added to select wells and fluorescence was monitored over time. Fevipiprant induced significant increases in apoptosis; this effect was abolished in CRTH2 KO neutrophils. Apoptosis was also higher in untreated CRTH2 KO neutrophils. Staurosporine was added as a positive control to stimulate apoptosis and zVAD as a negative control to inhibit apoptosis.

### CRTH2 deficiency attenuates lung injury in vivo

Having observed anti-inflammatory effects of CRTH2 antagonism on neutrophils in vitro, we predicted CRTH2 deficiency would ameliorate lung injury in vivo. To test this, we induced varying degrees of ALI by administering different doses of LPS. In line with our hypothesis, CRTH2 deletion resulted in improved mortality in severe ALI (10.0 μg/g LPS, ∼200μg/mouse) and attenuated weight loss and lung injury in mild ALI (0.25μg/g LPS, ∼5μg/mouse). Interestingly, this effect was only observed in male mice (Fig. 3). In line with these results, lung injury was improved in male CRTH2 KO mice with mild ALI at 96h.

**Figure 3:**
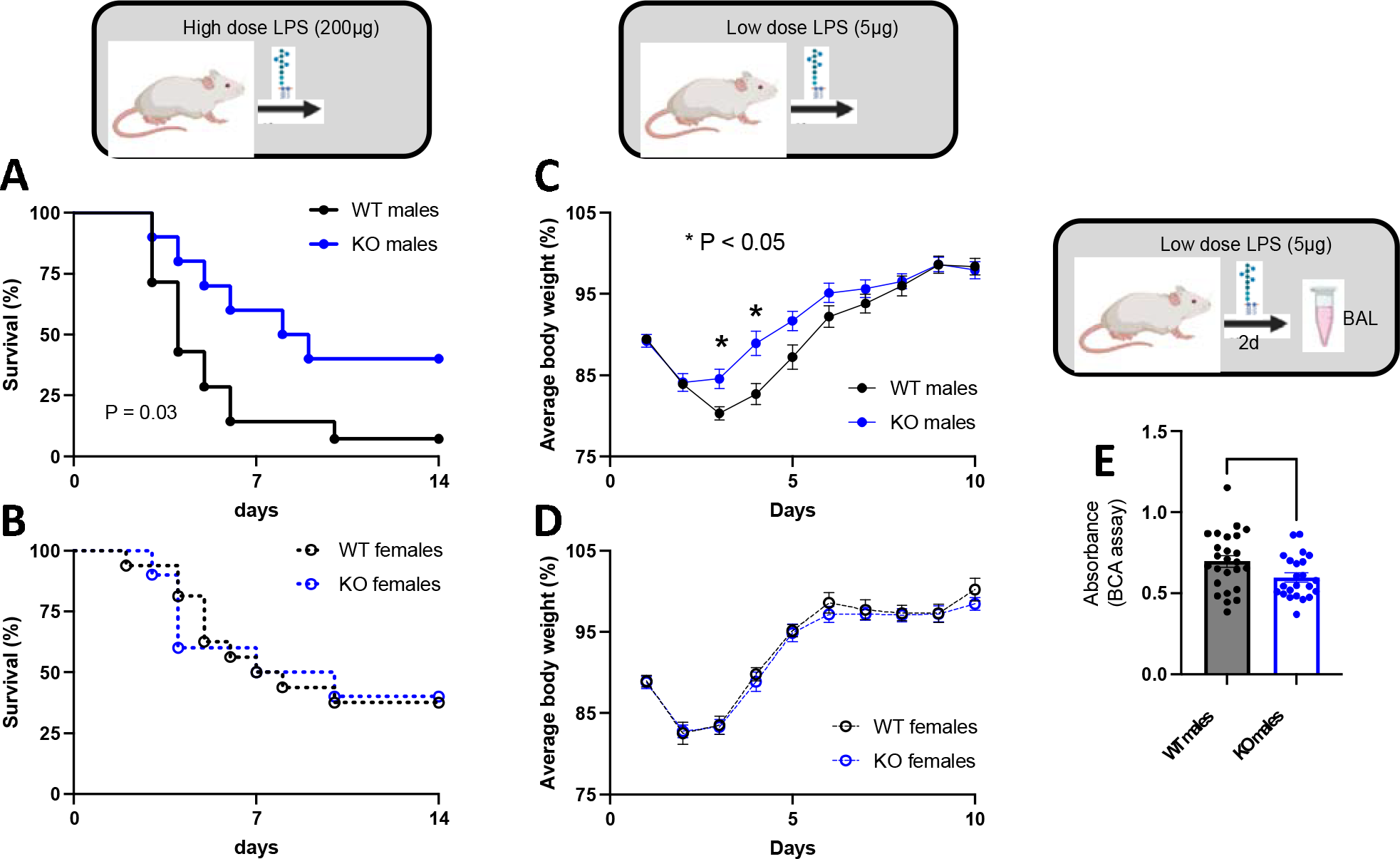
CRTH2 expression is protective during acute lung injury. Indicated doses of LPS were delivered to the lung to induce ALI. **A**, Survival was significantly increased in male CRTH2 KO mice, though not in females **(B). C**, Weight loss was significantly improved in male CRTH2 KO mice, though not in females **(D). E**, BAL total protein, measured via BCA assay, was decreased in male CRTH2 KO mice at the 96h timepoint in moderate ALI (15 μg of LPS).

### CRTH2 deficiency leads to heightened neutrophil effector function in vivo

Having observed improved outcomes after ALI in CRTH2 KO mice, we predicted that markers of neutrophilic inflammation would be decreased in these animals. However, we observed the opposite. Indeed, in both severe (Fig. 3A) and moderate ALI (Fig. 3B), we saw elevated levels of neutrophil elastase activity in CRTH2 KO mice.

### Transcriptomic analysis reveals exaggerated neutrophilic inflammation and suppression of Type 2 immunity in CRTH2 KO mice

To understand the apparent discrepancy between the pro-neutrophilic effects of CRTH2 in vitro and its anti-neutrophilic effects in vivo, we undertook bulk RNA sequencing (RNAseq) of lung tissue. As shown in Figure 4C, there were marked transcriptional differences in WT vs CRTH2 KO lungs. Consistent with our biochemical analyses (Fig. 4A-B), neutrophilic gene expression was highly upregulated in the CRTH2 KO (Fig. 4D). This was manifested by elevated expression of lymphocyte-derived cytokines (e.g. *il17a*), chemokines (e.g. *cxcl1*), and neutrophil effector proteins (e.g. *mmp8/9* and *camp*). We next conducted KEGG pathway mapping and GO enrichment analysis to identify candidate anti-inflammatory pathways that may be impaired in the CRTH2 KO mouse. This approach implicated decreased type 2 signaling – a finding that fits well with the established role of CRTH2 in type 2 immunity. Indeed, we observed downregulation of IL-4 and several downstream genes including *cldn2, muc5b, ptgs1* and *retnlg* – a central marker of alternatively-activated macrophages (AAM). A second AAM marker, *cxcl13*, was also decreased in CRTH2-deficient mice.

**Figure 4:**
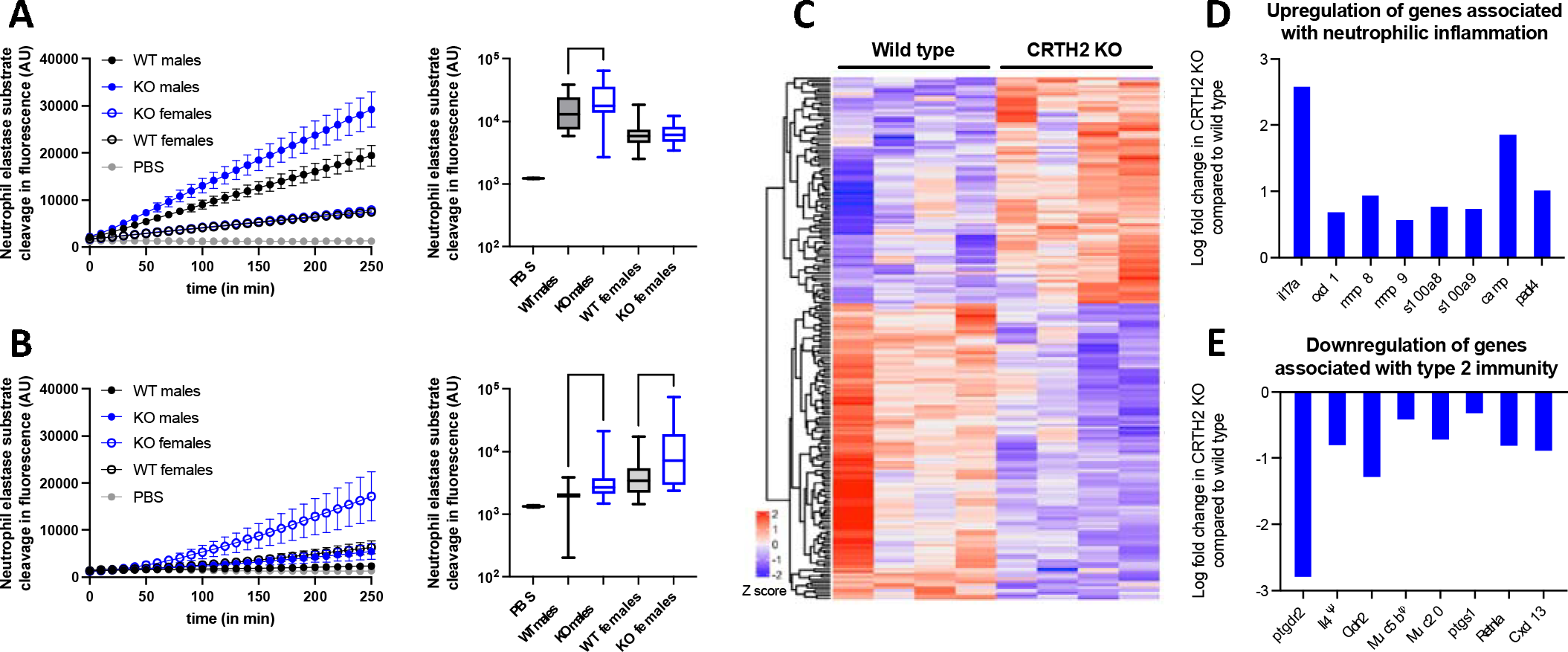
CRTH2 KO mice demonstrate exaggerated neutrophilic inflammation and defective type 2 immune responses. **A**, 200 μg of LPS was delivered to the lung to induce severe ALI, BAL was collected after 48h, and BAL fluid was assayed for neutrophil elastase activity. Data from the 200-minute timepoint is represented to the right. **B**, 15 μg of LPS was delivered to induce moderate ALI, BAL was collected after 10d, and BAL fluid was assayed for neutrophil elastase activity. Data from the 200-minute timepoint is represented to the right. **(C)**. 200 μg of LPS was delivered to induce severe ALI and bulk RNAseq was performed on lung tissue from male mice. A heat map of differentially expressed genes indicates transcriptional changes in CRTH2 KO mice. Genes associated with neutrophilic inflammation were significantly upregulated in CRTH2 KO mice **(D)**, while genes associated with type 2 immunity were downregulated **(E)** Differences were statistically significant, except where indicated by Ψ (p = 0.06) and ϕ (p = 0.10).

## Discussion

The role of CRTH2 in neutrophilic inflammation has not yet been studied in detail. This gap may be attributable in part to early reports demonstrating absent expression of CRTH2 on human blood neutrophils,^3^ leading investigators to focus on its function in type 2 immune cells such as eosinophils, which do express the receptor in blood. However, our results indicate that neutrophils do in fact express CRTH2, specifically after extravasation into the lung. Interestingly, one prior study similarly demonstrated tissue compartment-dependent expression of CRTH2; in patients with cystic fibrosis, neutrophils from blood were negative for the receptor, while sputum neutrophils stained positively.^29^ Thus, CRTH2 appears to be upregulated on neutrophils during extravasation into the lung in mice and humans. Further studies will be needed to evaluate if this phenomenon is generalizable to extra-pulmonary tissues as well.

Using a functional assay for apoptosis production we demonstrated that CRTH2 promotes cell survival in extravasated lung neutrophils. Neutrophils are known to synthesize and secrete PGD2, the endogenous ligand of CRTH2, at picomolar concentrations during swarming – a patterned response in which neutrophils seal off infected areas of tissue to prevent pathogen spread.^28^ In addition, DK-PGD2 has been shown to promote chemotaxis in bone marrow-derived murine neutrophils.^22^ Taking this literature precedent together with our findings, we propose the existence of a cell-autonomous PGD2-CRTH2 signaling mechanism that may promote neutrophil swarming and/or recruitment into the tissue. Indeed, leukotriene B4 – another neutrophil-derived eicosanoid – is known to mediate neutrophil recruitment via such an autocrine signaling during inflammation.^30^

Despite the direct pro-inflammatory effects on neutrophils, CRTH2 served a counterbalancing anti-neutrophilic role in vivo during ALI. Although our mechanistic studies are preliminary, they provide support for a model in which CRTH2 deletion leads to impaired type 2 immune signaling in the lung. This may be mediated by Th2 cells or type 2 innate lymphoid cells (ILC2s), which are the major sources of IL-4, IL-5, and IL-13. Our data specifically indicate a deficiency in IL-4, which acts on neutrophils via IL-4Ra to suppress their recruitment and activation.^31^

In summary, our findings indicate a dual role for CRTH2 during ALI: direct stimulation of inflammatory neutrophil responses, and indirect suppression of neutrophilic inflammation in vivo perhaps via IL-4. While these mechanisms would argue against CRTH2 antagonism as a therapeutic strategy in ARDS, it may prove useful for milder neutrophil-driven lung diseases such as chronic obstructive pulmonary disease, cystic fibrosis, and T2-low asthma. Further studies in relevant animal models will be necessary to test this hypothesis.

## Abbreviations

ALI: Acute lung injury
ARDS: Acute respiratory distress syndrome
CRTH2: Chemoattractant receptor-homologous molecule expressed on Th2 cells
ILC2: Type 2 innate lymphoid cells
KO: Knockout
LPS: Lipopolysaccharide
PGD2: Prostaglandin D2
RNAseq: Ribonucleic acid sequencing
WBC: White blood cells

